# Fluidic Programmable Gravi-maze Array for High Throughput Multiorgan Drug Testing

**DOI:** 10.1101/2025.06.18.660241

**Authors:** Henry C. Wong, Chris J. Collins, Joshna A. Jude Jose, Alyssa J. Villegas, Isha N. Bhakta, Andrew J. Collins, Gunjan Katara, Johar Kohana, Harpreet S. Saluja, John Collins

## Abstract

The high attrition rate of drug candidates in clinical trials underscores the urgent need for more predictive preclinical models that accurately replicate human physiology. Traditional 2D cell cultures and animal models often fail to predict human responses due to their limited physiological relevance, particularly for biologics and immunotherapies involving complex multicellular and cross-organ interactions. This highlights the need for modeling and measurements of multiorgan interactions at higher throughput, prompting the development of multiorgan-on-a-plate platforms. Here, we present OrganRX™, a modular, gravity-driven recirculation-based platform designed to imitate human organ function, physiological flow, immune cells circulation, and inter-organ communication in vitro. The Fluidic Programmable Gravi-maze Array (FPGA) technology integrates multiple organ models, including gut, liver, kidney, brain, tumor, and vascular compartments, within a microfluidic architecture designed to reproduce physiologically relevant shear stresses and gravity-driven recirculating flow that facilitates inter-organ communication. Using computational fluid dynamics (CFD) simulations and impedance-based flow validation, we confirmed accurate shear control across organ compartments. Organ-specific and multiorgan models were constructed using 3D extracellular matrix hydrogels and assessed for metabolism, toxicity, and senescence. Liver-kidney co-cultures demonstrated metabolic interplay via differential albumin and urea production. In addition, the platform was evaluated for biologics testing using immune-oncology models incorporating tumor spheroids, endothelial barriers, and circulating immune cells. Antigen-specific T-cells, checkpoint inhibitors, bispecific antibody and antibody-drug conjugate (ADC) studies demonstrated the ability to measure on-target tumor killing, off-target toxicity, cytokine release, and bystander effects across interconnected tissue compartments under dynamic recirculating conditions. The system enabled longitudinal evaluation of immune-mediated cytotoxicity, tissue-selective responses, and cross-organ signaling not readily captured in conventional static assays. Overall, the OrganRX™ platform offers a physiologically relevant, scalable, and automation-compatible platform for preclinical drug evaluation, biologics safety assessment, and disease modeling. Its ability to capture complex, dynamic inter-organ effects position it as a powerful tool for advancing translational research, mechanistic toxicology, and precision medicine.

## Introduction

The pharmaceutical industry faces significant challenges in advancing novel therapeutics from preclinical studies to clinical trials^1–4^, a critical stage in drug development, serves to evaluate the efficacy, safety, and pharmacokinetics of compounds before they can be tested in humans. Unfortunately, traditional preclinical models such as two-dimensional (2D) cell cultures and animal testing have consistently demonstrated poor predictive accuracy of human drug responses^5,6^. This has resulted in a staggering 90% failure rate for drug candidates in clinical trials, where unexpected toxicities and inadequate therapeutic efficacy become apparent only after considerable financial and time investments^7^. The limitations of 2D cell cultures stem from their inability to mimic the complexity of human tissue architecture and the microenvironment, leading to oversimplified models of cellular interactions and drug responses^8^. Animal models^9^, while offering some systemic insights, often fail to accurately replicate human physiology due to interspecies differences. This lack of translatability from animal models to human outcomes has contributed significantly to the inefficiency of the drug development pipeline, driving the need for more predictive, human-relevant preclinical models^10,11^.

To address these deficiencies, organ-on-a-chip (OOC) systems have emerged as promising tools to replicate human organ functions more accurately in vitro^12–16^. These platforms combine microfluidic technology with tissue engineering to create physiologically relevant models of human organ systems, enabling the study of complex biological processes^17–19^ such as drug metabolism, toxicity, and barrier functions. These systems offer distinct advantages over conventional models, including the ability to simulate dynamic fluid flow, shear stress, and nutrient transport, all of which are critical for maintaining the viability and functionality of human tissues in vitro^20,21^.

However, despite their promise, current OOC technologies still face significant limitations for studying systemic drug effects. A major challenge lies in recreating the intricate physical and biochemical cues that govern cellular behavior in vivo. Accurate simulation of fluid dynamics, particularly the shear stress and pressure conditions experienced by cells in different organs, remains technically challenging due to the need for precise control over microfluidic flows and pressure gradients^22–24^. Building on the OOC foundational advancements, multiorgan-on-a-chip (MOOC) platforms were developed to address the need for studying systemic drug effects. MOOC systems aim to replicate the complex interactions between different organs involved in processes such as drug absorption, distribution, metabolism, and excretion (ADME)^25^. These platforms often incorporate liver, kidney, heart, and lung models to evaluate drug candidates’ effects across multiple organ systems simultaneously^26^. While these systems have demonstrated significant potential in providing a more holistic view of drug effects, their widespread application in preclinical drug testing remains limited by the challenges discussed earlier, including scalability, robustness, and the accurate simulation of human physiological conditions^27^. Moreover, the scaling of MOOC platforms for high-throughput drug screening applications remains an unresolved issue. While some platforms have demonstrated the ability to model single organs with reasonable accuracy, the integration of multiple organ systems into a unified, scalable platform has proven difficult^28^. Each organ has unique physiological needs, and replicating the microenvironments necessary to sustain multiple organ types within the same system presents substantial technical hurdles^29^. Furthermore, many existing MOOC platforms lack the robustness and reproducibility required for widespread adoption in pharmaceutical R&D, with significant variability in results across experiments^30^ allowing for the controlled delivery of nutrients, gases, and other factors necessary for cell survival^31^. Further, proper fluid flow is critical for simulating blood circulation, which governs nutrient and drug delivery as well as waste removal^32,33^. The fluid shear stress on the endothelial lining of blood vessels differs across organs^34^, in vivo and significantly influences cellular behavior, gene expression, and drug metabolism. Replicating these dynamic forces in vitro, while maintaining the appropriate flow rates and pressure gradients across multiple organ models, is technically challenging due to the precise engineering required to avoid flow inconsistencies or mechanical disruptions^35^. Furthermore, scaling up multiorgan platforms using external pumps, valves, and tubing may not be practical or user-friendly for preclinical pharmacological studies, limiting their broader adoption^36^.

Specifically, the increasing complexity of modern immunotherapies necessitates preclinical platforms capable of modeling interactions between tumors, immune cells, vascular barriers, and healthy tissues within a connected physiological environment^37^. Therapeutic modalities such as checkpoint inhibitors^38^, T-cell engagers^39^, bispecific antibodies^40^, ADCs^41,42^, and engineered cell therapies exert their effects through dynamic immune-mediated mechanisms that extend beyond a single organ compartment. Consequently, both therapeutic efficacy and adverse events often emerge from systemic immune responses involving multiple tissues. Multiorgan immune-on-a-chip systems^43^ provide the ability to investigate immune-cell circulation, tissue-specific trafficking, cytokine propagation, on-target and off-target cytotoxicity, bystander effects and immune-mediated organ toxicities in real time. By integrating tumor, liver, brain, gut, kidney, and vascular compartments under recirculating flow, these platforms could mimic human physiology and offer a potential tool for predicting clinical responses, identifying safety liabilities, and accelerating the development of safer and more effective immunotherapies^44^.

To address the limitations of current MOOC platforms, we have developed the Fluidic Programmable Gravi-maze Array (FPGA)—a modular, gravity-driven multiorgan platform that robustly simulates human physiological conditions through unidirectional flow guided by a precisely engineered maze architecture. This multiple repeated units of MOOC in a plate format, termed *OrganRX™*, integrates organ-specific models—liver, kidney, lung, gut, and brain—within a microfluidic circuit designed to replicate key aspects of inter-organ communication, fluid dynamics, and biochemical signaling. OrganRX™ features organ-specific microenvironments and dynamic flow to simulate complex pharmacokinetic interactions in a controlled, customizable manner. The system offers tunable flow rates and pressure gradients, allowing tailored preclinical testing to meet diverse experimental needs. The gravity-driven recirculation system mimics physiological blood flow and shear stress across organ compartments to potentially become a human on a plate (HOAP). Furthermore, it will enable accurate modeling of drug distribution, metabolism, and inter-organ responses, with real-time monitoring of drug concentrations, metabolites, and cellular activity^45^. The OrganRX™ platform has the potential to support future clinical trials-on-a-chip^46^ by enabling the integration of multiple human tissue models that capture aspects of drug metabolism^47^, and inter-organ interactions^48^ within a patient-relevant system^49^.

In this study, we employed computational and experimental tools to demonstrate the generation of net-unidirectional recirculating flow and physiologically relevant shear stresses within the OrganRX™ platform. Using a set of biological and biochemical assays, we further demonstrated the platform’s ability to support organ-specific functions, inter-organ communication, and immune cell circulation for evaluating multiple biologics for immunotherapy.

## Methods and Materials

### Development of Multiorgan Plate and Recirculation System

The OrganRX™ platform consists of two recirculation circuits separated by a blood-brain barrier (BBB) compartment, as illustrated in Fig. 1a. The brain model is positioned within the inner recirculation circuit, where lower flow rates and shear stresses are maintained to better reflect the physiological microenvironment of neural tissues. Brain cells may be cultured either as two-dimensional monolayers within the wells or as three-dimensional tissue constructs within the inner triple-cross channels. Other organ models, including gut, liver, kidney, and vascularized BBB compartments, are connected to the outer recirculation circuit. The epithelial tissue compartments are dimensioned according to physiologically relevant functional requirements and intended inter-organ interactions, whereas the endothelial recirculation channels are designed to provide shear stress levels comparable to those encountered in the microvasculature. Unless a BBB is established, all organ compartments share a common circulating medium to facilitate biochemical communication and systemic signaling. A distinguishing feature of the OrganRX™ design is its gravity-driven recirculation architecture, which promotes net-unidirectional transport of soluble factors, therapeutic agents, and immune cells throughout the interconnected organ compartments. Additional flow-rectification elements, such as Tesla valves or diffuser-nozzle structures, may be incorporated to further enhance directional transport. Although transient flow reversal may occur during portions of the tilting cycle, the resulting directional changes are of substantially lower magnitude and duration than the forward flow phase, resulting in minimal effects on cell alignment, barrier integrity, mechano-transduction, cytokine signaling, and tissue maturation.

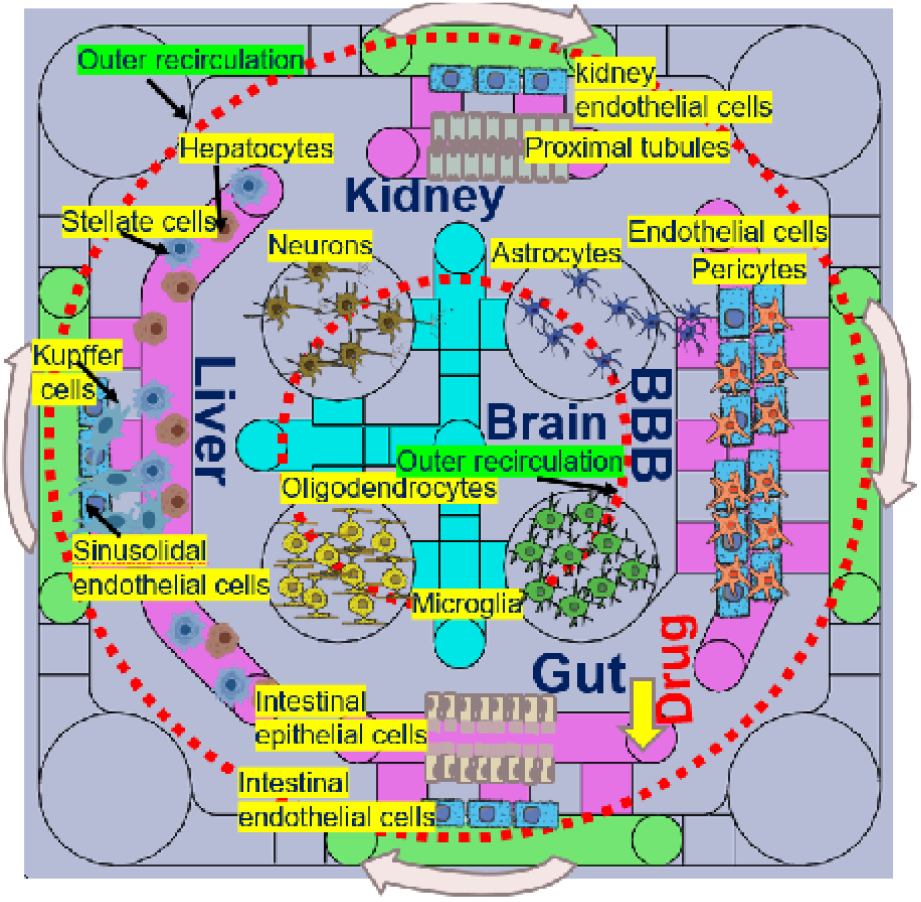

**Fig. 1.**
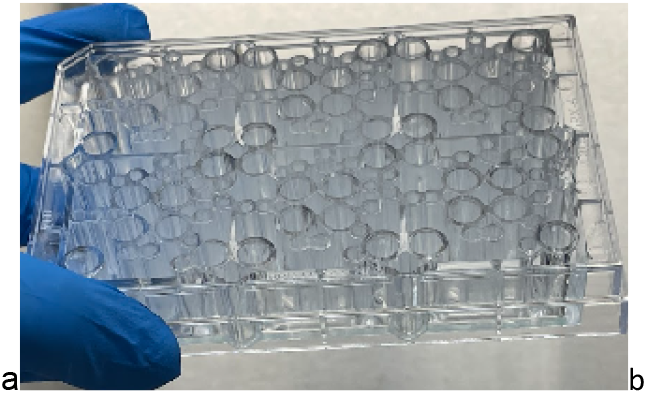
(a) Schematic of multiorgan plate, OrganRX™ (b) Injection-molded glass bottom multiorgan OrganRX™plate with 6-multiorgan units

The multiorgan plate with six repeated units was injection molded with polystyrene and bonded with a 0.17mm glass at the bottom as shown in Fig. 1b. The width of the channels is 2mm and depths ranging from 0.2mm at the bridges to 0.7mm at the main endothelial and epithelial channels. The epithelial channels are loaded first with the gel and they stop at the bridges due to Laplace stop force. The plate is compatible with standard well format and is accessible to imaging systems and multiplate readers. The plate is also compatible with automated liquid dispensing robots such as Opentrons, OT2. Gravity-driven recirculation is established by tilting the plate to drive flow between four inner corner wells and four outer corner wells, maintaining net-unidirectional flow in the inner and outer fluidic loops.

The OrganRX™ plate operates under controlled recirculation with physiologically relevant shear flow using the OrganRX™ system. Following sterilization with 70% ethanol, the system is placed inside a standard cell culture incubator. To demonstrate high-throughput capability, we developed an expandable system capable of cascading up to 30 plates (arranged in a 2×3×5 configuration) across the x, y, and z axes. Notably, two such units can fit within a conventional incubator, enabling parallel experimentation of 720 multiorgan models. Additionally, we developed and published a Bluetooth-enabled iOS app on the App Store, which provides remote control of the recirculation system. This app enables users to select from multiple shear flow rates, set programmable waiting times between tilt-direction changes, and reset the system as needed—enhancing both precision and usability.

### Experimental Setup to study Net-Unidirectional Flow

We investigated the fluid dynamics^50^ and shear stress within microchannels of OrganRX™ multiorgan-on-a-plate system under different experimental conditions. We used impedance measurements to verify the flow rate of 1M NaCl salt solution injected into a corner well of the organ plate. An impedance spectrometer from Zurich Instruments was connected to a computer running dedicated impedance data acquisition software, which was used to capture and record impedance changes of the salt solution during transient flow condition in real time. The injection is adjusted so that the initial velocity is negligibly low. The impedance was measured from the well of the delivery of the salt solution and an adjacent well of the organ plate to determine the average time between the two impedance peaks during the gravity-driven flow. A set of stainless-steel electrodes were inserted into the wells of the organ plate, with one probe positioned in Channel 1 (red, Fig. 2) and the other in Channel 2 (blue, Fig. 2), corresponding to a known distance within the microfluidic channel. From the time of travel, the velocity of the flow and shear stresses were calculated based on the time intervals between impedance peaks.

**Fig. 2.**
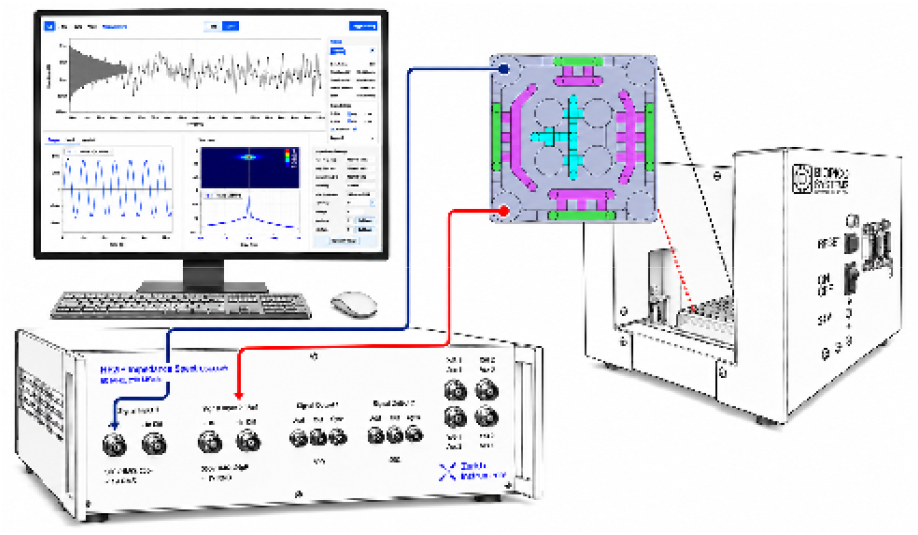
Experimental Setup for Shear-flow Rate Measurements using high speed impedance acquisition

The organ plate was subjected to controlled tilting at various z height increment steps. To control the flow dynamics of the OrganRX™, an iOS App was employed to set eight predefined tilt increments. Trial experiments were conducted to examine how fluid flow and shear stress were influenced by both the fluid volume and the addition of solutes.

By analyzing the time intervals between impedance peaks at the two probe locations, we calculated the fluid velocity (*v* ) between them. The Length (= 27*mm*) between the two probes, and the velocity was obtained by dividing this distance by the average time interval (<Δt>) between the impedance peaks:

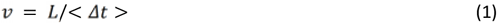

### Calculation of Shear Stresses in the Organ Plate

The schematic in Fig. 3 shows the organ plate placed in the fixture of the organ system. Tilting of the plate to various angles is accomplished by moving its four corners using four stepper motors in the organ system simultaneously. As the motors move, if an extension point’s incremental height moves to the highest position, *z*_*l =*_ *Δh* _*system*_, the diagonally opposite connection point is moved to the lowest position *z*_3_= 0. The other two connection points are moved to the half-height position 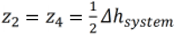 . This causes the fixture to be tilted at an incremental tilt angle, which is defined as:

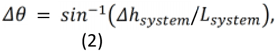

**Fig. 3.**
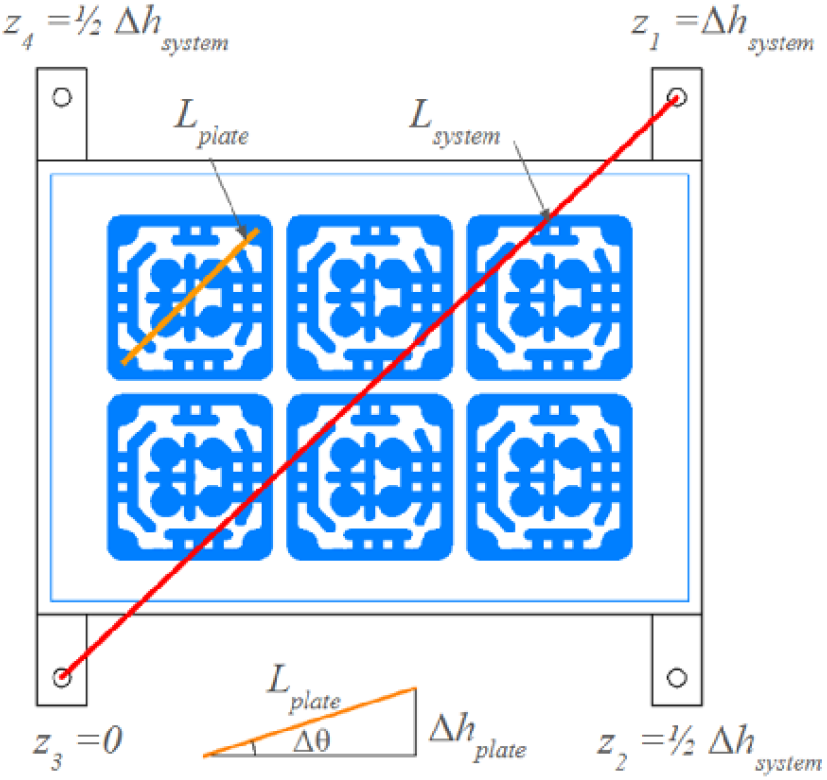
Schematic of the organ plate in the tilting fixture illustrating the definition of the incremental tilt angle and incremental height.

Where *L* _*system*_ = 189 mm is the distance between the extensions in the OrganRX™ system. This incremental tilt causes a height difference in the media levels in the plate. The incremental height in one unit of the organ plate is the height difference between the well at the highest position and the well at the lowest position. This is given by:

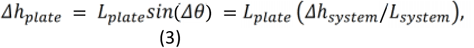

Where *L*_*plate*_= 27 mm is the distance between corner wells in one unit in the organ plate. The pressure gradient of this height difference of the fluid in one unit is given by:

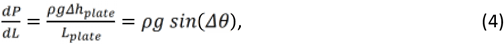

Where*ρ* = 1 g/cm^3^ is the density of the water modeled as media and *g* =9.8 m/s^2^ is the gravitational acceleration.

Because flow within the channels reaches steady state much more rapidly than media transport between the interconnected wells, a quasi-steady-state assumption was applied. Under this assumption, velocity magnitudes and wall shear stresses were calculated to represent the initial or maximum values generated immediately following tilting of the organ plate. The equations describing the velocity and shear stress fields, as well as the average flow speed, mean wall shear stress, and volumetric flow rate, are presented in the Supplemental section. Over time, the height difference between media levels decreases as fluid moves from the highest to the lowest well. This behavior can be captured using a mass balance across the interconnected wells, resulting in an exponential decay of the incremental height for the plate:

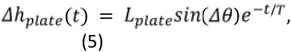

where the time constant is:

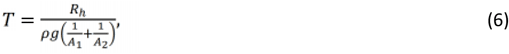

*R*_*h*_ is the hydraulic resistance of the rectangular channel, *A*_1_and *A*_2_are the cross-sectional areas of the wells ^51,52^. Since the wells have the same diameter *D*= 6 mm, this reduces to

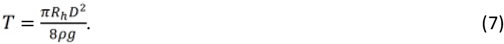

For our system, this evaluates to *T* = 0.85 seconds.

### Computational Fluid Dynamics (CFD) Simulations

Precise control and prediction of fluid flow are critical where physiologically relevant shear stresses must be recreated to support organ-specific function. Traditional analytical models often fail to capture the complex, transient behaviors arising from gravity-driven flow in non-planar geometries. To ensure that the OrganRX™ platform accurately mimics in vivo-like fluid dynamics across multiple organ compartments, computational fluid dynamics (CFD) simulations were employed to model media movement during plate tilting. We used Simcenter STAR-CCM+ to perform CFD simulations of transient fluid flow within the OrganRX™ plate during gravitational tilting. The volume fraction module was utilized, assigning the 2 mL of medium in the organ plate physical properties corresponding to the culture media, while the surrounding region was defined as air. No-slip boundary conditions were applied to all channel walls, and zero pressure was imposed at the media surface in each well. The initial condition assumed an equal media height across all wells. To simulate a high-shear condition, the plate was tilted by 25°, elevating the lower right corner relative to the upper left. This induced a pressure gradient across the system, generating fluid flow.

### Cell Expansion in T-Flasks

All cell lines used in the organ model studies were expanded in T-75 flasks (Corning, Cat. #430641U) according to the manufacturer’s recommended protocols for thawing and subculture. For the brain organ model, SK-N-SH neuroblastoma cells (ATCC, Cat. #HTB-11) and HMC3 human microglia (ATCC, Cat. #CRL-3304) were used. For the gut–liver–kidney model, cell lines sourced from ATCC—HepG2 (Cat. #HB-8065), HK-2 (Cat. #CRL-2190), and T84 (Cat. #CCL-248)—were utilized. The cells were expanded under recommended growth conditions by ATCC, in EMEM or DMEM supplemented with 10% FBS and 1% Penicillin-Streptomycin. For the vascular model, HUVEC (Lonza, Cat. #: C2519AS) and Normal Human Lung Fibroblasts (NHLF) (Lonza, Cat. #: CC-2512) were cultured in Endothelial Growth Medium-2, prepared by adding Endothelial Growth Medium-2 SingleQuots Supplements (Lonza, Cat. #CC-4176) to Endothelial Basal Medium-2 (Lonza, Cat. # CC-3156) according to the manufacturer’s instructions, and DMEM/F-12 media supplemented with 10% FBS and 1% Penicillin-Streptomycin, respectively. All cultures were maintained in a humidified incubator at 37 °C with 5% CO_2_, and passaged at 70–90% confluence prior to use in 3D encapsulation or seeding within OrganRX™ plates.

### CD8+ T-cell Culture and Expansion

HLA matched and mismatched CD8+ T-cells (Ignyte Bio, Cat. #TC-03) were thawed according to manufacturer’s instructions and seeded in T-75 flasks in RPMI media with glutamax (Sigma-Aldrich, Cat. #R8758) and supplemented with 10% FBS and 1% Penicillin-Streptomycin, and 1 ng/mL recombinant human Interleukin-2 (ScienCell, Cat. #101-02). Prior to experiments, CD8+ T-cells were activated with 2 ug/mL Concanavalin A (Sigma-Aldrich, Cat. #C2010) for 24 hours. Antigen-specific CD8^+^ T cells (Ignyte Bio, Cat. #TC-13) were thawed according to manufacturer’s instructions and seeding in the OrganRX™ plate in RPMI media with glutamax and supplemented with 10% FBS and 1% Penicillin-Streptomycin, and 1 ng/mL recombinant human Interleukin-2.

### Spheroid Formation

The AggreWell400 (StemCell Technologies, Cat. #34411) plate was prepared by adding 500 uL of Anti-Adherence Rinsing Solution (StemCell Technologies, Cat #07010) to each well and centrifuging at 300 x g for 5 minutes. The anti-adherence rinsing solution was discarded. For 3-D tumor modeling, 2,000,000 T84, HMC, SKN cells/well were plated in DMEM or EMEM media as previously prepared. HLA matched human Hepatocytes (Lonza, Cat. #HUCPG) and Kupffer cells (Lonza, Cat. #HLKC-200K) were plated at 208,000/well and 225,000/well respectively in Spheroid Formation Medium (Hepatocyte Culture Medium, 20% FBS, and 25 mM HEPES). Hepatocyte Culture Medium (HCM) was prepared by adding Hepatocyte Culture Medium SingleQuots Kit (Lonza, Cat. #CC-4182) to Hepatocyte Basal Medium (Lonza, Cat. #CC-3199) according to the manufacturer’s instructions. Immediately after seeding the cells, the plate was centrifuged at 100 x g for 5 minutes and placed in a humidified incubator at 37 °C with 5% CO_2_ for 5 days to allow for mature spheroid formation. After 5 days, 50% media changes were performed using DMEM, EMEM, or Hepatocyte Culture Medium.

### 3-D Organ Model Experiments

3D organ models were embedded in extracellular matrix (ECM) hydrogels tailored to support specific tissue types. Initial 3-D culture experiments were performed using rat tail type I collagen gel (Corning, Cat. #354236) at a final concentration of 4 mg/mL with pH-adjusted to 7.4 using NaHCO_3_. The collagen ECM consisted of rat tail type I collagen gel, NaHCO_3_ and HEPES solution combined in a 8:1:1 ratio. Fibrinogen from bovine plasma (Sigma-Aldrich, Cat. #F8630) at a final concentration of 3 mg/mL was dissolved in sterile PBS and mixed 1:1 with thrombin from bovine plasma solution (Sigma-Aldrich, Cat. #T4648) at 2 U/mL prepared in heparin-supplemented media to form a fibrin hydrogel^53^. This formulation was used for encapsulating HUVECs, and NHLFs. For the brain organ model, a 1:1 co-culture of SK-N-SH neuroblastoma cells and HMC3 human microglia was seeded at a density of 30,000 cells/µL per unit. For the gut–liver–kidney model, cell lines sourced from ATCC such as HepG2, HK-2, and T84 were seeded along with HuVEC in 1:1 ratio. All experiments were performed using the OrganRX™ plate, with tissue channel gel volumes ranging from 15 µL (kidney), 20 µL (gut) and 50 µL (liver). Each co-culture suspension was combined with a fibrinogen/thrombin-based ECM and incubated at 37 °C for 40 minutes in a humidified 5% CO_2_ incubator to allow for gelation. OrganRX™ recirculation system was configured to achieve organ-specific shear stress levels, delivering approximately 0.5 dyn/cm^2^ for epithelial models and 4 dyn/cm^2^ for endothelial barrier models.

### Image-Based Assays

Organ models cultured in the OrganRX™ plate were monitored daily using brightfield microscopy (NyOne, SynenTec) to assess cell morphology, attachment, and tissue integrity. Whole-plate imaging was performed at 4× magnification, while higher-resolution imaging of selected organ units was conducted at 20× magnification. For each selected field, 20 Z-stack images were acquired spanning a depth range of 0.2 mm to 0.7 mm to capture the three-dimensional architecture of the tissue construct. Fluorescent imaging was used to visualize labeled cells within the organ system. Brain microvascular endothelial cells expressing green fluorescent protein (GFP) were imaged using a standard FITC filter set (excitation: ∼488 nm; emission: ∼520 nm). All imaging was performed using an automated inverted fluorescence microscope equipped with motorized stage control and Z-stack capability. Image data were subsequently processed for qualitative and quantitative assessment of cellular distribution, barrier coverage, and structural organization.

### Establishment of a Vascular Model

To develop a functional vascular model^54^ within the OrganRX™ platform, a fibrinogen/thrombin-based ECM was injected into the designated vascular channel (volume 25 uL) (as shown previously in Fig. 1a with label BBB) using a micropipette. The gel was allowed to polymerize at 37 °C in a humidified incubator with 5% CO_2_ for 30–45 minutes. To avoid premature gelation, it is recommended to mix the thrombin gel with cells and fibrinogen for each unit, as shown in Fig. 4 with the sequence of the steps in mixing. Following gelation, 2 ml media is added into the wells and placed in the recirculation system. To develop functional in vitro models^55–57^ within the OrganRX™ platform, a type I collagen hydrogel was also used. The collagen ECM consisted of 5 mg/mL rat tail type I collagen gel (Corning, Cat. #354236), NaHCO_3_ and HEPES solution combined in an 8:1:1 ratio. The collagen matrix was allowed to polymerize at 37 °C in a humidified incubator with 5% CO_2_ for 30–45 minutes.

**Fig. 4.**
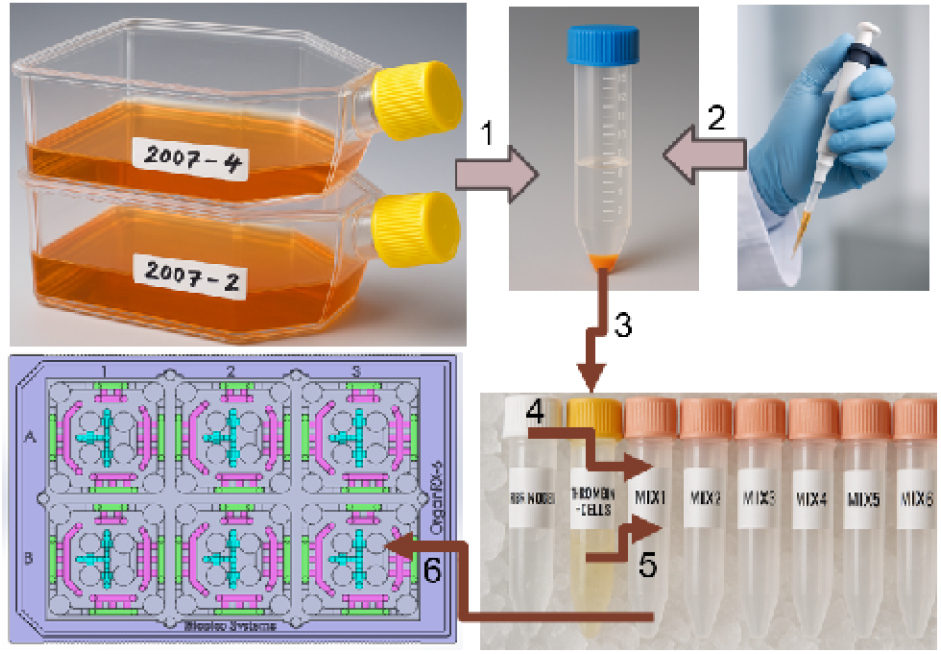
3-D gel with cells for seeding into the OrganRX™ plate

### Cell Painting using JUMP Cell Painting Kit

For Cell Painting^58^, cells the organ model is grown for 5 days under flow conditions. Prior to staining, models were washed once with pre-warmed 1× PBS and fixed using 3.7% paraformaldehyde for 15 minutes at room temperature. After fixation, cells were permeabilized with 0.1% Triton X-100 in PBS for 10 minutes and subsequently washed twice with PBS. The staining protocol followed the manufacturer’s instructions provided in the JUMP Cell Painting Kit (Revvity, Cat. #CPJ-001), which includes six fluorescent probes targeting distinct cellular compartments: Hoechst 33342 (nuclei), SYTO 14 (cytoplasm), Concanavalin A–Alexa Fluor 488 (endoplasmic reticulum), Wheat Germ Agglutinin–Alexa Fluor 555 (plasma membrane and Golgi), Phalloidin–Alexa Fluor 568 (F-actin), and MitoTracker Deep Red (mitochondria). Each dye was diluted in a staining buffer to the recommended working concentrations and applied simultaneously for 2 hours at room temperature, protected from light. Following incubation, cells were washed three times with PBS to remove excess dye, and plates were imaged using the LSM 780 confocal microscope from Carl Zeiss at UC Irvine (Optical Biology Core) using linear unmixing^59^. at appropriate excitation/emission settings for each fluorophore. In this linear unmixing, spatial expression patterns of Angiotensin-Converting Enzymes^60^ (ACE1 and ACE2) in the model are computationally distinguished using multiple fluorescent signals with overlapping spectra.

### Preparation of Aβ_42_ Oligomers (Aβ_42_Os)

Aβ_42_ oligomers (Aβ_42_Os) were prepared following previously published^61^ protocols with minor modifications. Synthetic Aβ_42_ peptides (GenScript, Cat. #RP10001) were first pretreated with hexafluoroisopropanol (HFIP) to monomerize the peptides and then air-dried. The dried peptide film was resuspended in dimethyl sulfoxide (DMSO) (Sigma-Aldrich, Cat. #D8418) to generate a 5 mM stock solution. This stock was further diluted to a final concentration of 100 µM in phenol red-free DMEM/F-12 medium (Thermo Fisher Scientific, Cat. #11039021). The solution was vortexed for 15 seconds and incubated at 4 °C for 24 hours to allow oligomer formation. The resulting Aβ_42_O solution was used directly in experiments or aliquoted and stored at –80 °C for single use to minimize freeze– thaw variability.

### Reactive Oxygen Species (ROS) Assay

Reactive oxygen species (ROS) generation was assessed^62^ in co-culture models following exposure to metabolic stress. Cells were pretreated with the drugs (for eg. D/10Q) in media for 2 days, followed by treatment with 0.1 µM doxorubicin (Sigma-Aldrich, Cat. #D1515) for 24 hours. To quantify intracellular ROS levels indicative of cellular senescence^63^, the fluorescent probe 2′,7′-dichlorodihydrofluorescein diacetate (H_2_DCF-DA) (Thermo Fisher Scientific, Cat. #D399) was added to the culture medium within OrganRX™ units at a final concentration of 10 µM and circulated on the OrganRX perfusion platform for 4 hours at 37 °C. Following incubation, the organ models were imaged on 4X and 20X using a fluorescence microscope with an excitation wavelength of 480 nm and an emission wavelength of 540 nm. Fluorescent images were analyzed using intensity as a readout for ROS accumulation across the co-culture system.

### Measurement of Senescence-Associated β-Galactosidase Activity

Senescence-associated β-galactosidase (SA-β-gal) activity was assessed^64^ in fixed co-culture organ models using a commercial SA- β-gal staining kit (Cell Signaling Technology, Cat. #9860), following the manufacturer’s instructions. The induce senescence, the organ models cultured within the OrganRX plates were treated for 48 hours with either Aβ_42_ oligomers (Aβ_42_O) or doxorubicin (DOX). Following fixation, samples were stained for SA-β-gal activity. Quantitative measurement was performed by reading absorbance at 420 nm and 600 nm using a Synergy2 microplate reader (Agilent). Additionally, stained organ cultures were imaged using light microscopy to qualitatively assess senescent cell morphology and distribution.

### Functional Biochemical Assays

Biochemical assays^65^ were performed using an Agilent Synergy2 microplate reader configured with a modified XML file to accommodate sporadic 1536-well formats. Assays employing absorbance, fluorescence, or chemiluminescence detection were conducted according to manufacturer protocols. Urea levels in conditioned media were quantified using a colorimetric assay kit (BioAssay Systems, Cat. #DIUR-100), and albumin was measured using a quantitative ELISA (Bethyl Laboratories, Cat. #E88-129). Cytochrome P450 1A2 (CYP1A2) enzymatic activity was assessed using a luciferin-based chemiluminescent assay (Promega, P450-Glo™ CYP1A2 Assay, Cat. #V8751), following chemical induction of the organ models. Standard curves and controls were included for all assays to ensure quantitative accuracy.

### Sample Recovery and Gene Expression Analysis

To retrieve samples for gene expression analysis, OrganRX™ plates were first cleaned with 70% ethanol, and a one-sided adhesive tape (3M, Cat. #465) was applied to the bottom surface. Adhesion was secured using a handheld roller (VWR, Cat. #89097-614). Plates were then frozen at –86 °C for approximately 2 hours until the bottom glass was loose enough to be separated. Following freezing, the adhesive tape and 0.17 mm cover glass were carefully removed. Gel samples were scraped into 1.5 mL RNase-free microcentrifuge tubes (Eppendorf, Cat. #022431081) for downstream RNA isolation.

For efficient gel collection, media-only channels were scraped first to allow partial thawing (∼30 seconds), followed by recovery of cell-containing gel sections. In cases where gel melted into the perfusion channels, a long pipette tip (Rainin, Cat. #17014079) was used to gently push the gel through. Optionally, dowel pins (McMaster-Carr, Cat. #91595A005) were used to block channels and prevent gel leakage.

Total RNA was extracted^66^ using the Direct-zol RNA Microprep Kit (Zymo Research, Cat. #R2062) according to the manufacturer’s protocol. Complementary DNA (cDNA) was synthesized from isolated RNA using the SuperScript™ VILO™ cDNA Synthesis Kit (Invitrogen, Cat. #11754050).

Quantitative real-time PCR (qRT-PCR) was performed using Maxima SYBR Green/Fluorescein qPCR Master Mix (2X) (Thermo Scientific, Cat. #K0241) on a QuantStudio™ 3 Real-Time PCR System (Applied Biosystems, Cat. #A28567). Gene-specific primers were ordered from Integrated DNA Technologies (IDT) and used as follows: p21 : Forward: 5′-CGA TGG AAC TTC GAC TTT GTC A-3′; Reverse: 5′-GCA CAA GGG TAC AAG ACA GTG-3′ and PAI-1 : Forward: 5′-ACC GCA ACG TGG TTT TCT CA-3′; Reverse: 5′-TTG AAT CCC ATA GCT GCT TGA AT-3′

Data collection and analysis were conducted using QuantStudio™ Design and Analysis Software (Applied Biosystems, Cat. #A28219). Fluorescence data were exported in .xlsx format, and quantitative assessment was based on the multicomponent plot for SYBR signal (Cycles vs. Fluorescence).

### Dasatinib and Quercetin Dosing

Dasatinib and Quercetin are both senolytics^67,68^ that target senescent cells, thereby preventing the senescent induction of surrounding cells by lowering senescence-associated secretory phenotype (SASP) factors secretion. After culturing the cells with 7 days of maturation in the incubator, stock solutions of 500 uM Dasatinib and 15 mM Quercetin (D / 10Q) in DMSO were used to prepare 6 working solutions of varying concentrations in media. Working solutions were added to the 6 units of the OrganRX plate. The following working concentrations of D / 10Q were prepared: 0 µM (0.7% DMSO), 0.25/2.5 µM, 0.5/5 µM, 1/10 µM, 2/20 µM, 3/30 µM. Doxorubicin (0.1 µM) was used to induce senescence within the cells in the organ plate.

### Biologics Dosing

In order to model tumor killing, the bispecific antibody cibisatamab (IchorBio, Cat. #ICH5041) was evaluated at concentrations 0, 0.003, 0.03, 0.3, 1, and 3 ug/mL. The antibody-drug conjugate Labetuzumab govitecan (GLPBIO, Cat. #GC74567-1) was also evaluated at concentrations 0, 0.45, 1.5, 4.5, 15, 45 ug/mL. In order to model human liver-immune toxicity, the CTLA-4 immune checkpoint inhibitor ipilimumab (Syd Labs, Cat. #C050P) was evaluated at 0.0845, 0.169, 0.4225, 0.845, and 1.69 mg/mL and compared to control small molecule troglitazone (AGScientific, Cat.# T-2469) at concentrations 0, 0.0565, 0.02825, 0.01415, 0.00565, and 0.002825 mg/mL.

### Establishment of Liver-Immune model

To establish a functional liver immune model within the OrganRX™ platform, 400 Hepatocyte+Kupffer (H+K) spheroids were embedded in fibrin gel and seeded in the OrganRX™ plate. The gel was allowed to polymerize at 37 °C in a humidified incubator with 5% CO_2_ for 30 minutes before addition of 1 mL HCM media is added into the wells and placed in the recirculation system. Matched or mismatched CD8+ T-cells were added in suspension in 1 mL to each unit of the OrganRX™ plate with an effector to target ratio of 1:1. The OrganRX™ plate was placed in a 37 °C in a humidified incubator with 5% CO_2_ in the recirculation system. In parallel, the above process was repeated for a 6-well plate. After addition of matched or mismatched CD8+ T-cells, the 6-well plate was placed 37 °C in a humidified incubator with 5% CO_2_ in the recirculation system under static conditions.

### 3-D Tumor Killing in Multiorgan Model

CellTrack Green CMFDA (ABP Biosciences, Cat. #C039) was dissolved in DMSO to 10 mM. HMC3 cells were incubated for 30 minutes at 37 °C in Hank’s Balanced Salt Solution (HBSS) and 10 mM HEPES with 20 uM CellTrack Green CMFDA. CytoTrace™ Red CMTPX (AAT Bioquest, Cat. #22015) was dissolved in DMSO to 2 mM. CD8+ T cells were incubated for 30 minutes at 37 °C in HBSS and 20 mM HEPES with 15 uM CytoTrace™ Red CMTPX. Both cells were centrifuged and the solution was removed and replaced with the cell’s respective media.

### Statistics

All studies were conducted in triplicate. Bar plots represent the mean ± SEM. Statistical analyses were performed using Excel, MATLAB, or custom Python scripts. Error bars are omitted for fluidic experiments involving high-frequency data acquisition. For image-based ROS measurements, outliers were removed by sorting pixel intensities in ascending order and excluding the top and bottom 10% of values. For phenylbutyrate and curcumin sequential dosing experiment images, the drug treatment images are normalized with control (no drug treatment) images.

## Results and discussion

### Net-Unidirectional flow in OrganRX™ endothelial channel

In a shear flow experiment, 2 mL of DI water was introduced into the well, and 1 M NaCl solution was incrementally dispensed into two opposite side channels in 40 steps (20 steps in each channel) of 0.05 mL each corresponding to 360° angle of rotation of the organ plate. For each step of NaCl dispensed, impedance measurements were taken, allowing us to track the corresponding peaks. In these impedance measurements, a constant current of 1mA is sent to the device under test and instantaneous voltages are measured. The graph presented in Fig 5a is an example dataset used to demonstrate the calculation of delta time (Δt) between impedance peaks at two different probe locations (Channel 1 and Channel 2). These are real voltage signals obtained from the impedance spectroscopy system during fluid flow in the microfluidic channels. To calculate the Δt between peaks, we identified a kink in the graph for both Channel 1 and Channel 2. We calculated the time difference between kinks at the two probe locations.

**Figure 5.**
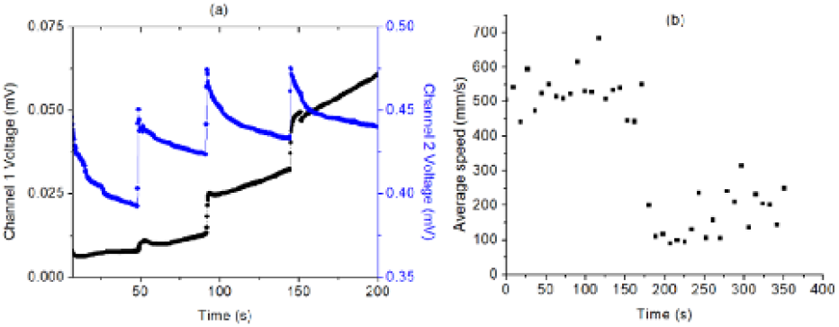
(a) Real voltage measurements from the impedance spectroscope during the NaCl injections (b) Forward flow in the organ plate is represented from revolution angle of 0 to 180° while reverse flow is represented from 180° to 360°

Using the time difference and the distance between probe locations, we calculated the speed for each of the 40 steps. The speeds recorded over a full rotation of the organ plate in the recirculation system are plotted in Fig. 5b. The first 20 data points, corresponding to angles 0° to 180°, represent the forward flow, while the second 20 data points, corresponding to 180° to 360°, represent the reverse flow. As the forward flow velocities are significantly higher than those of the reverse flow, the system effectively operates in a semi-unidirectional flow regime. While manual dispensing using a syringe introduced minor variations in the volume and consequently velocity due to differences in applied force for injection, the velocities at each peak were expected to remain largely consistent.

### Shear Stresses in the Organ Plate

The CFD analysis enabled dynamic mapping of the pressure and velocity magnitude fields within the microchannels of the OrganRX™ plate. Fig. 6 shows plots of the pressure and velocity distributions after the organ plate is tilted at 25° for 300 ms. The pressure head is highest at the bottom of the well that experienced the most vertical displacement from the tilting. Maximum flow is observed at the outer endothelial channel (88 to 176 mm/s) and lower flow (< 80 mm/s) is observed in the epithelial channels.

**Figure 6.**
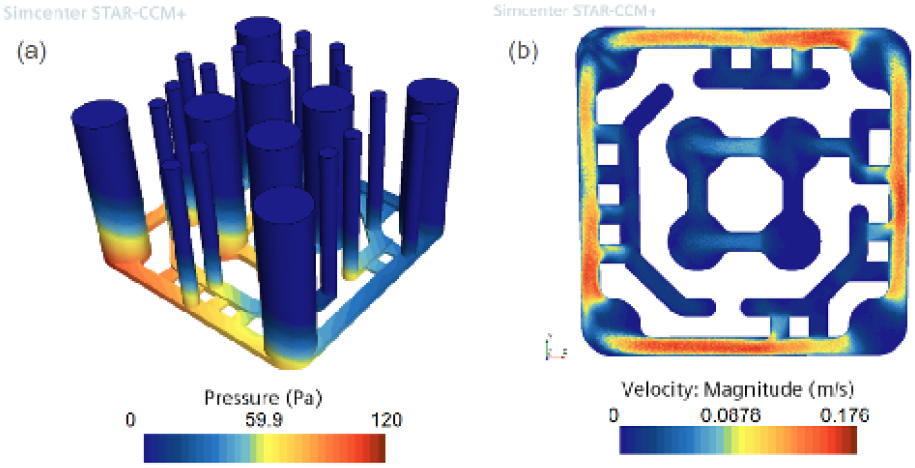
CFD results of organ plate tilted at 25° for 300 ms showing the (a) pressure field and (b) velocity magnitude

Figure 7a shows a plot of the decay of the maximum speed in the endothelial channel over time. The CFD takes into account the delay in media flow as it takes about 118 ms for the media to reach its maximum speed of 214 mm/s after the initial tilt. This was not modeled with the theoretical calculations. We also see that the maximum velocity magnitude is reduced by 90% after approximately 2 seconds. This slow decay from the CFD results matches the results predicted by the theoretical calculations where we modeled the incremental height as an exponential decay. This validates the assumptions that we made earlier that the time scale for fluid flow between the interconnected wells is larger than the time scale of flow in the microchannels. Thus, it is valid to use the steady-state values for velocity magnitude and shear stress as the maximum values.

**Figure 7.**
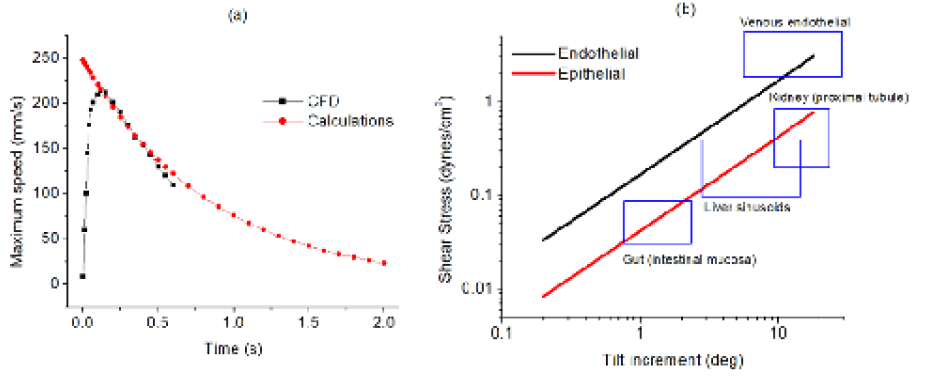
(a) Plot of the computed average velocity magnitude from the CFD and theoretical calculations as a function of time (b) Average wall shear stress as a function of the organ system tilt increment for endothelial and epithelial regions. Also plotted are the physiological range of shear stress for venous endothelial cells, kidney (proximal tubule), liver (sinusoids), and gut (intestinal mucosa).

**Fig. 8.**
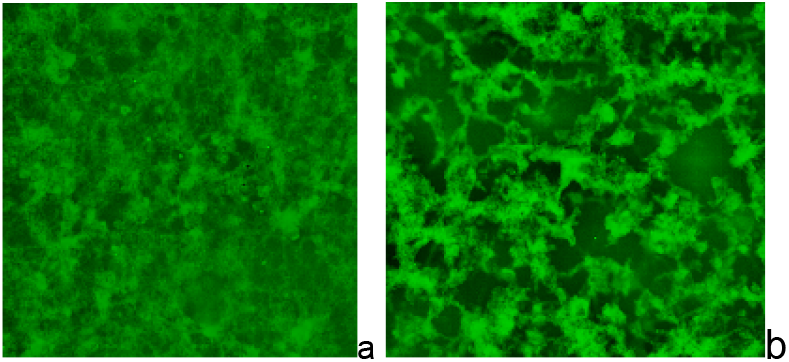
Vascular structure formation with endothelial cells (HUVECs) (a) without fibroblast cells (b) with fibroblast cells.

From the CFD data, we were also able to estimate the relative velocity magnitudes between the endothelial and epithelial regions. Using Equations 4, 5, 7 and 8, we calculated the average wall shear stress in the outer endothelial channel of the organ plate as a function of the tilt increment. The ratio of velocity magnitudes from the CFD simulations were then used to scale the shear stress results for the epithelial regions. Fig. 7b shows that the average wall shear stresses for these regions increases linearly with the tilt increment of the organ plate. We also plotted the range of shear stress values for venous endothelial cells^69^, kidney (proximal tubules)^70,71^, liver (sinusoids)^72,73^, and gut (intestinal mucosa)^74,75^. Flow from the endothelial channel into the brain compartment is regulated by the blood-brain barrier region, resulting in only low levels of physiologically relevant interstitial flow within the brain tissue region. This confirms that the OrganRX™ System is able to provide physiologically relevant shear stresses for endothelial and epithelial cells by controlling the tilting angle.

### Development of a Vascularized Endothelium

To establish a vascularized endothelial network^53,54,76^, Human HUVECs at 1.25 million cells and LHLFs at 625,000 cells at a 2:1 ratio were co-cultured within a fibrin-based 3D matrix in the blood-brain barrier vascular channel region of the OrganRX™ plate. 3D Matrix Composition is Fibrinogen: 6 mg/mL and Thrombin: 4 U/m. The organ plate was placed on the continuous recirculation OrganRX™ system and maintained under standard culture conditions (37 °C, 5% CO_2_). Vascular structure formation was monitored by brightfield and fluorescence microscopy, with visible network formation observed between days 2 and 7 post-seeding. The vascular structure without and with fibroblasts after 5 days of recirculation is presented in Fig.8.HuVECs alone may correspond to large-vessel venous endothelium whereas HuVECs with fibroblasts perfused microvascular/capillary-like vascular network model.

In another experiment, HUVECs were used to form perfusable 3D vascular structures, which were subsequently stained using the Cell Painting JUMP Kit. High-resolution spectral unmixing imaging was performed using confocal microscopy to characterize the structural and morphological integrity of the vascularized network as shown in Fig. 9. The image shows how HUVECs can self-organize into vessel-like tubes with central lumens (empty spaces) expressing ACE enzymes on their surfaces. The vascularized HUVECs with expression of ACE1 and ACE2 could reveal important biological insights especially in the context of vascular function, inflammation, or disease modeling. ACE1 upregulation often correlates with inflammation and leakiness. ACE2 may be reduced or redistributed during such stress, as seen in viral infection or oxidative damage.

**Fig. 9.**
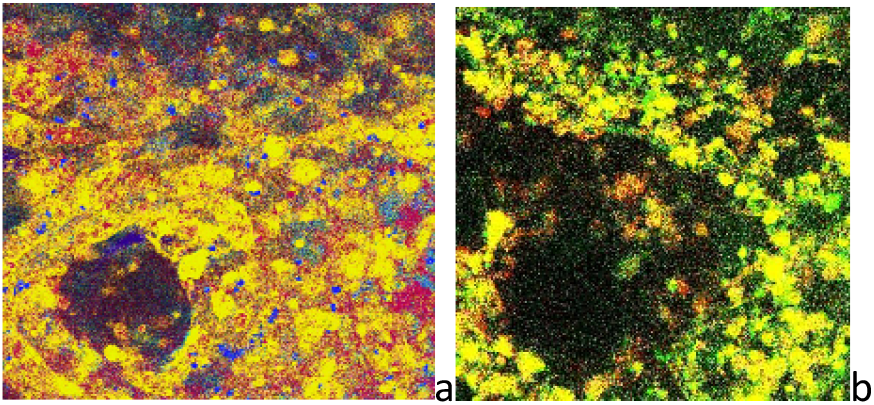
(a) Vascular HuVEC cells using cell painting using spectral unmixing Angiotensin-Converting Enzyme 1 (b) Angiotensin-Converting Enzyme 2

The vascularized HUVECs with expression of ACE1 and ACE2 could reveal important biological insights especially in the context of vascular function, inflammation, or disease modeling. ACE1 upregulation often correlates with inflammation and leakiness. ACE2 may be reduced or redistributed during such stress, as seen in viral infection or oxidative damage.

### Albumin and Urea Generation from Multiorgan Models

Initial multi-organ experiments, which included gut, brain, liver, and kidney models simultaneously, led to complex and inconsistent results that were difficult to interpret and reproduce. Subsequent studies focused on simplified dual-organ systems^77^, such as liver– kidney, to better investigate inter-organ communication under controlled conditions. Later, kidney and liver organ models were established in a 3D co-culture system incorporating HUVECs to emulate vascular integration. The multiorgan constructs consisted of 1 M HUVECs co-cultured with 1 M HT-2 cells and 1 M HepG2 cells respectively to model renal tissue and hepatic tissue. Functional output was assessed by quantifying albumin and urea concentrations in the culture medium (Fig. 10). By day 8, albumin levels in the liver– kidney co-culture were reduced relative to the liver-alone condition, whereas urea levels were elevated in the multiorgan model compared to the hepatic monoculture, suggesting dynamic interorgan metabolic interactions^70^.

**Fig. 10.**
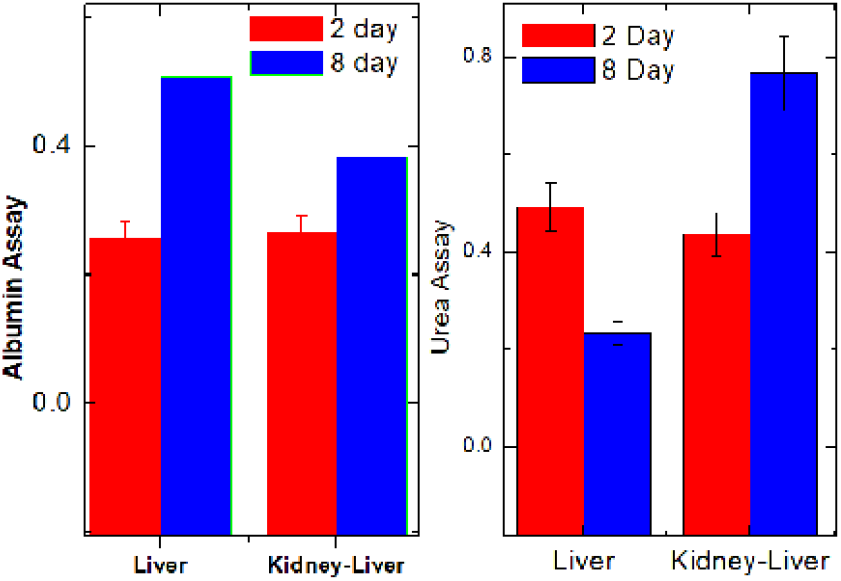
Albumin and urea evaluation in the liver–kidney co-culture model on the 2nd day and 8th day

The observed reduction in albumin levels within the liver–kidney with HUVEC co-culture systems may be attributed to several factors^78,79^. First, in a multiorgan configuration, albumin secreted by HepG2 hepatocytes is likely distributed across a larger extracellular volume and may be partially absorbed or retained by non-hepatic cells. This dilution or adsorption effect can result in lower measurable concentrations of albumin in the collected media. Second, both HT-2 and HuVEC cells possess surface receptors and endocytic pathways capable of albumin binding or uptake, thereby reducing its extracellular concentration without affecting overall hepatic synthesis. Additionally, inter-organ crosstalk may modulate hepatic function directly. Co-culture with HT-2 or HUVEC cells may influence HepG2 metabolic activity through nutrient competition, the secretion of inflammatory cytokines, or shear stress exposure in the presence of recirculation. These factors may suppress albumin synthesis or secretion over time. In contrast, urea concentrations were elevated in the multiorgan system compared to liver monocultures, a result consistent with increased nitrogenous metabolic load. The presence of HT-2 and HUVEC cells likely introduces additional amino acid turnover and metabolic waste, thereby providing more ammonia substrates for hepatic urea cycle activity and stimulating urea synthesis by HepG2 cells. Moreover, in this model, renal urea clearance mechanisms are absent, as HT-2 cells do not possess the specialized transport and excretory functions of proximal tubule epithelial cells.

### Evaluation of Brain Disease Model and Neuroprotection

To induce cellular senescence, we established a 3D brain model using a 1:1 co-culture of HMC3 microglia and SK-N-SH neuroblastoma cells, seeded at a total cell density of 30,000 cells/µL. Senescence was triggered by treatment with hydrogen peroxide (H_2_O_2_)^80^, doxorubicin^81^, or Aβ oligomers^82^. After 7 days of culture, 10 µL of the H_2_DCF-DA probe was added to each unit and incubated for 4 hours on the OrganRX™ circulation system for reactive oxygen species (ROS) analysis. ROS fluorescence imaging was performed using 20× magnification, for quantification. H_2_O_2_ was applied at varying concentrations to induce oxidative stress, and the resulting ROS levels were measured. In a separate set of experiments, the combination of dasatinib and quercetin (D+10Q) was administered at different concentrations to evaluate its neuroprotective potential. There was a concentration dependent increase in ROS levels following H_2_O_2_ exposure, as seen in Fig. 11. D+10Q treatment resulted in a reduction in ROS, indicating a protective effect against oxidative stress.

**Fig. 11.**
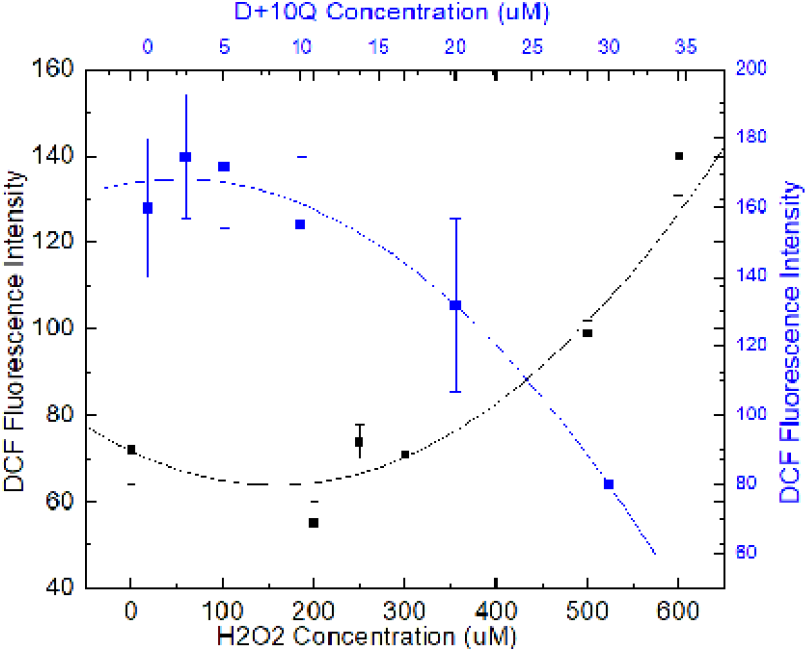
ROS assay after stressing using H2O2 [left] and ROS assay after protecting using D+Q drugs [right])

We observed that ROS imaging in the brain model varies depending on the type of stressor applied. Representative ROS images following exposure to H_2_O_2_ and DOX are shown in Fig. 12a and Fig. 12b.

**Fig. 12.**
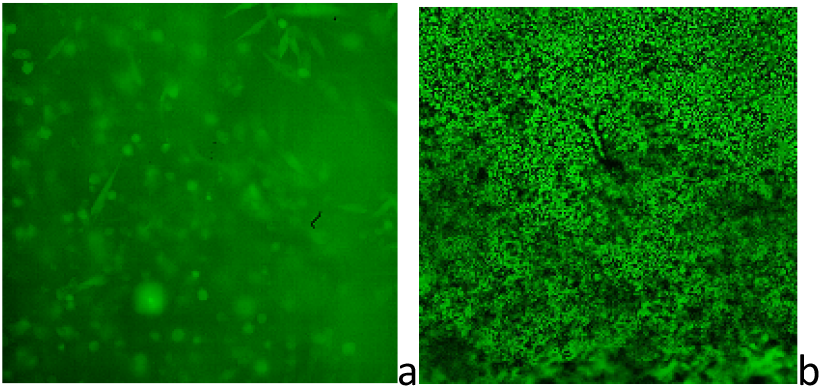
(a) ROS image after stressors using H2O2 (b) ROS assay after stressors using DOX

With relevance to Alzheimer’s disease drug screening, we used Aβ oligomers to challenge the brain organ model and verified the senescence experienced by the organ model using RTPCR and SAβGal assay. The neuroprotective effects of D+Q drugs are exhibited through SAβGal assay and RTPCR with genes p21 and PAI-1 as presented in Fig. 13.

**Fig. 13.**
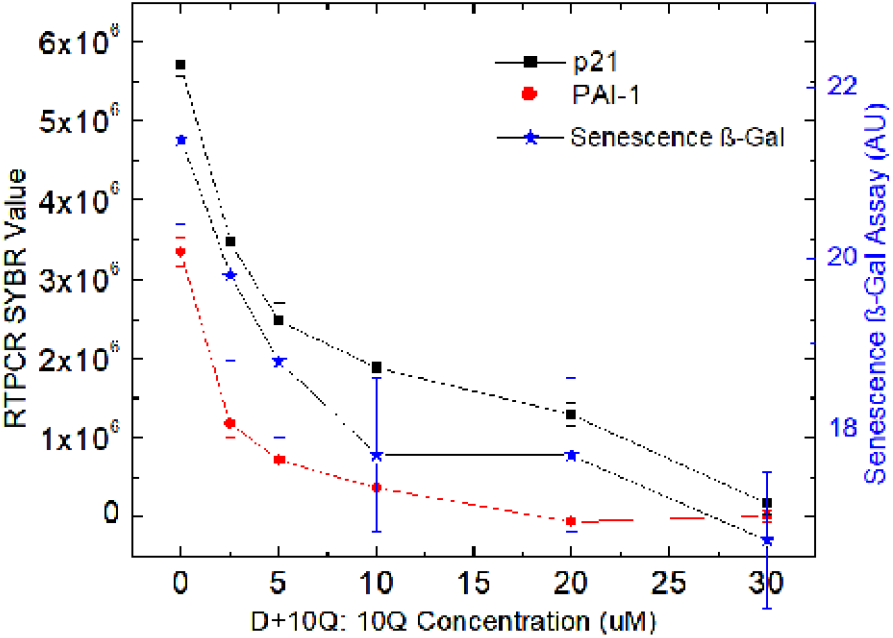
RTPCR assay (left axis) and Senescence β-Gal assay after stressing using Aβ oligomers followed by neuroprotection using D+Q drugs.

### Human Liver-Immune Modeling for Immunotoxicity

To establish a human liver-immune model, we first compared the OrganRX™ platform under flow conditions to a static 6-well plate. Both plates contained HLA matched human hepatocyte and kupffer spheroids embedded in a fibrinogen/thrombin-based ECM and HLA matched or mismatched CD8+ T-cells in suspension. At various timepoints, both plates were analyzed for cytokines IFN-y, IL-6, IL-17A, and TNF-a and LDH release (Fig.14a). The OrganRX™ platform shows sustained cytokine production when compared with static culture systems as evident at later time points (168-336h). This suggests that dynamic recirculating flow supports prolonged immune-cell viability and signaling. The static 6-well system likely experiences nutrient depletion, uneven cytokine distribution, and reduced cell-cell interaction over time. The OrganRX™ platform demonstrated enhanced cytotoxicity, as evidenced by earlier and greater LDH release compared with the static 6-well system. The flow-based system also showed clearer separation between matched and mismatched conditions, indicating more effective immune-mediated killing and improved effector-target interactions under dynamic recirculating flow. In contrast, the static 6-well culture exhibited delayed and reduced LDH responses with less distinct differentiation between experimental groups. The continuous recirculating flow in the OrganRX™ system enhances cytokine distribution, nutrient transport, and cell-cell communication, resulting in stronger and more sustained T-cell activation and cytotoxicity compared with static culture, demonstrating improved assay sensitivity and physiological relevance for immuno-oncology and toxicity studies. Using the model, we studied toxicity of a checkpoint inhibitor, ipilumumab and a positive control small molecule troglitazone which is known for drug-induced liver toxicity, as shown in Fig. 14b.

**Fig. 14.**
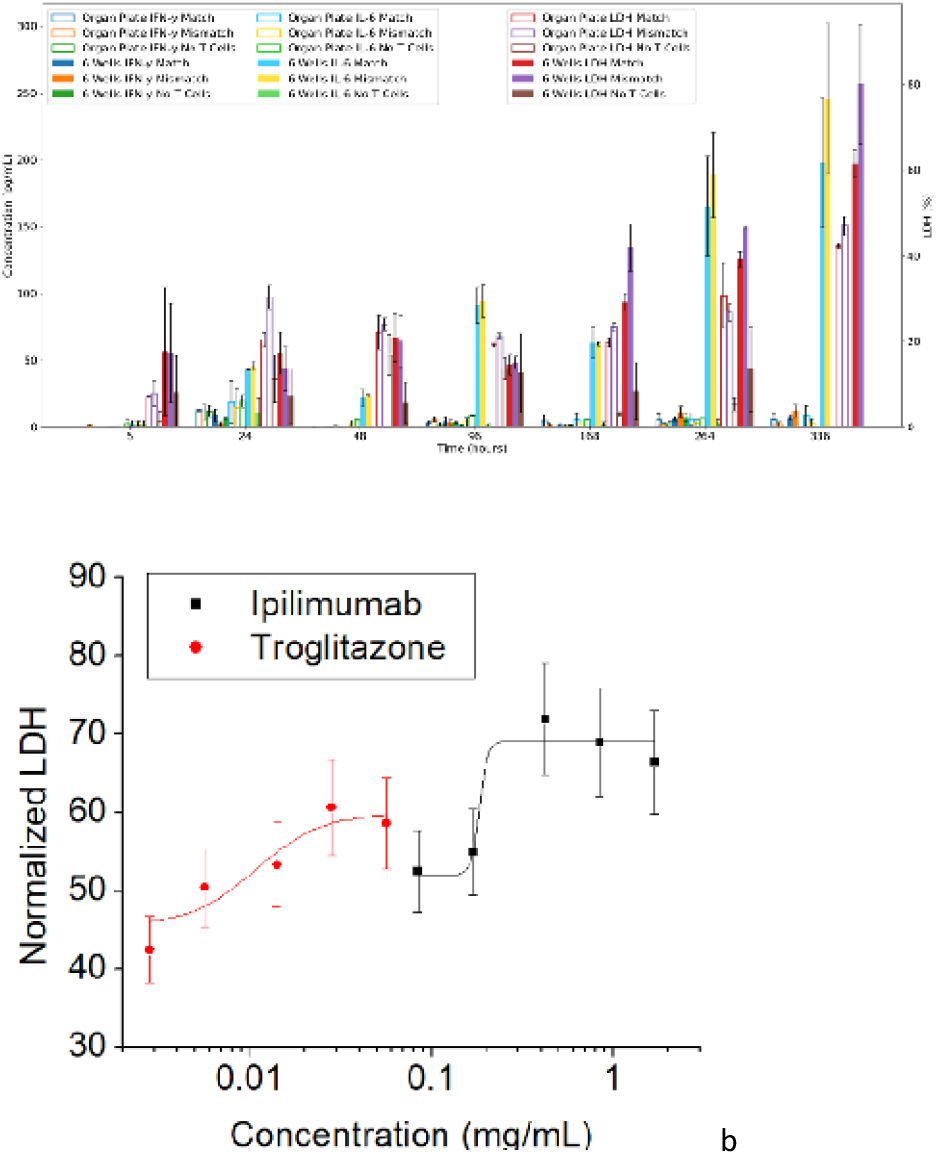
(a) Comparison of cytokines and LDH released from human HLA matched and mismatched human primary hepatocytes and Kupffer cell spheroid models with CD8+ T cells at different time points comparing OrganRX™ flow and 6-well under static conditions. (b)Toxicity profile of Ipilimumab and control molecule troglitazone.

### Modeling Immunotherapies using Multiorgan Models

#### Tumor Killing with Antigen-specific CD8^+^ T cells

Antigen-specific CD8^+^ T cells were co-cultured with on-target CEA^+^ gut tumor spheroids (E:T=1:1) and off-targets neuroblastoma and microglia spheroids in multiple channels to evaluate tumor-targeting efficiency. Tumor spheroids expressing the target antigen (CEA) exhibited clear dose-dependent cytotoxicity, while off-target tissues showed significantly reduced direct killing. Importantly, the system captured dynamic killing kinetics, reflecting sustained immune engagement under recirculating flow conditions. The imaging viability data as shown in Fig. 15a demonstrate selective cytotoxicity toward gut spheroids over time following exposure to antigen-specific CD8^+^ T cells. Gut spheroids viability progressively decreased, while neuroblastoma and microglia spheroids maintained relatively high viability, indicating preferential on-target immune-mediated killing with limited off-target toxicity. The schematic in Fig. 15b illustrates the interconnected recirculating organ configuration, enabling simultaneous evaluation of on-target tumor killing and off-target tissue effects under dynamic flow conditions. These results demonstrate the capability of the OrganRX™ platform to evaluate both efficacy and safety of immunotherapies in a physiologically relevant multi-organ human tissue environment. The graph in Fig. 15c demonstrates the time-dependent response of on-target and off-target tissues to antigen-specific CD8^+^ T-cell therapy within the OrganRX™ multi-organ platform. On-target (gut spheroids) viability progressively decreased over time, indicating effective immune-mediated tumor killing, while off-targets (microglia and neuroblastoma spheroids) maintained high viability, suggesting minimal killing effects.

**Fig. 15.**
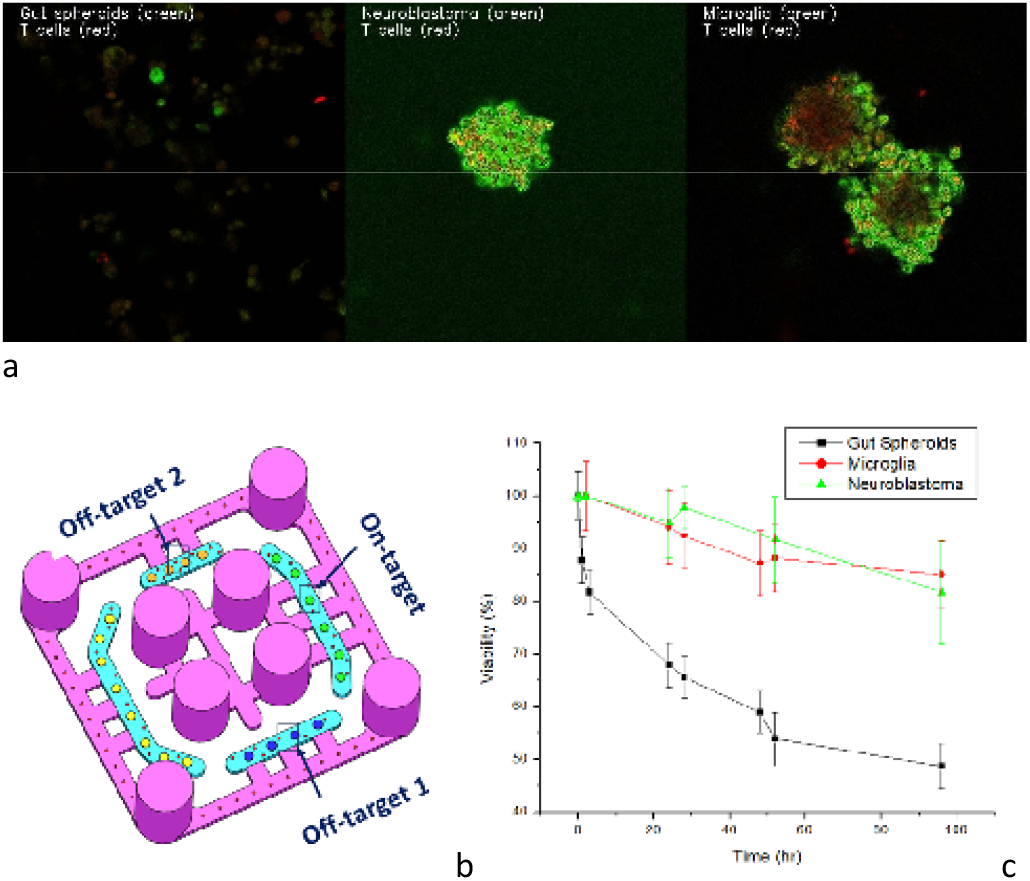
(a) Representative image and (b) Configuration for on-target / off-target effects (c) evaluation of tumor killing kinetics using fluorescent imaging for viability in a physiologic immune environment.

### Tumor Killing with Bispecific Antibodies

The OrganRX™ platform was used to evaluate the bispecific antibody, cibisatamab in the same interconnected multi-organ system with gut spheroids as on-target together with off-target neuroblastoma spheroids and microglia under recirculating flow conditions. To demonstrate efficient tumor cell killing and immune-mediated cytotoxicity, CD8+ T cells with E:T=1:1 are introduced from the corner wells. The treatment induced a time- and dose-dependent reduction in gut spheroids viability, demonstrating effective on-target immune-mediated tumor killing by antigen-specific T cells, while microglia and neuroblastoma remained relatively stable throughout the study, indicating limited off-target toxicity as shown in Fig. 16a. A dose-dependent reduction in gut spheroid viability, recorded after 120 hours is presented in Fig. 16b. As in Fig. 16c, the OrganRX™ platform was configured to model bystander effects by integrating on-target and off-target tissues within an interconnected recirculating system. This configuration enabled simultaneous evaluation of direct tumor killing and unintended effects on neighboring healthy tissues under physiologically relevant immune communication and soluble factor exchange. Tumor killing kinetics evaluated in Fig. 16d using longitudinal fluorescent viability imaging within the platform demonstrated progressive reduction in gut spheroids as well as off-target tissues viability over time, enabling real-time assessment of therapeutic efficacy and tissue-specific toxicity.

**Fig. 16.**
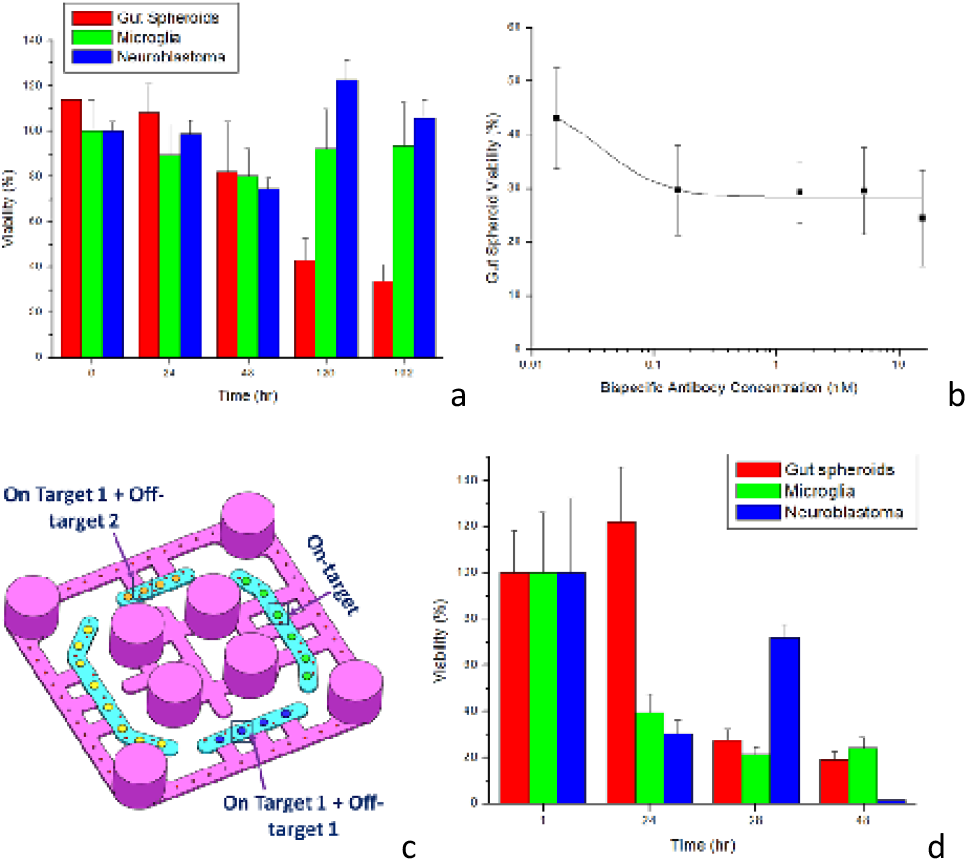
(a) Evaluation of on/off-target tumor killing kinetics of c Bispecific antibody, cibisatamab (b) Effect of bispecific antibody concentration on target cells’ viability after 120 hours of treatment. (c)Configuration for Bystander Effects (d) evaluation of tumor killing kinetics using fluorescent imaging for viability in a physiologic immune environment.

### Tumor Killing with Antibody-Drug Conjugate

Antibody-Drug Conjugate, Labetuzumab Govitecan targets CEA-expressing tumors with a topoisomerase-inhibiting payload in the presence of CD8+ T cells E:T=1:1. The evaluation of on-target and off-target killing kinetics following treatment with the ADC in the multi-organ platform is shown in Fig. 17a. Gut spheroids showed progressive reductions in viability over time, demonstrating effective ADC-mediated tumor killing, while neuroblastoma spheroids and microglia maintained relatively higher viability, indicating reduced off-target toxicity. The effect of the ADC concentration on target cell viability after 120 hours of treatment is shown in Fig. 17b. Increasing ADC concentrations resulted in a dose-dependent decrease in target cell viability, demonstrating concentration-dependent therapeutic efficacy with saturation at higher doses. The evaluation of on-target and off-target killing kinetics using bystander effects under interconnected co-culture conditions is shown in Fig. 17c. The OrganRX™ platform enabled simultaneous assessment of direct tumor killing and indirect toxicity to neighboring tissues, revealing differential viability responses among gut spheroids, microglia, and neuroblastoma within a physiologically relevant recirculating microenvironment. The model reveals both on-target tumor killing and bystander toxicity caused by payload diffusion and immune amplification.

**Fig. 17.**
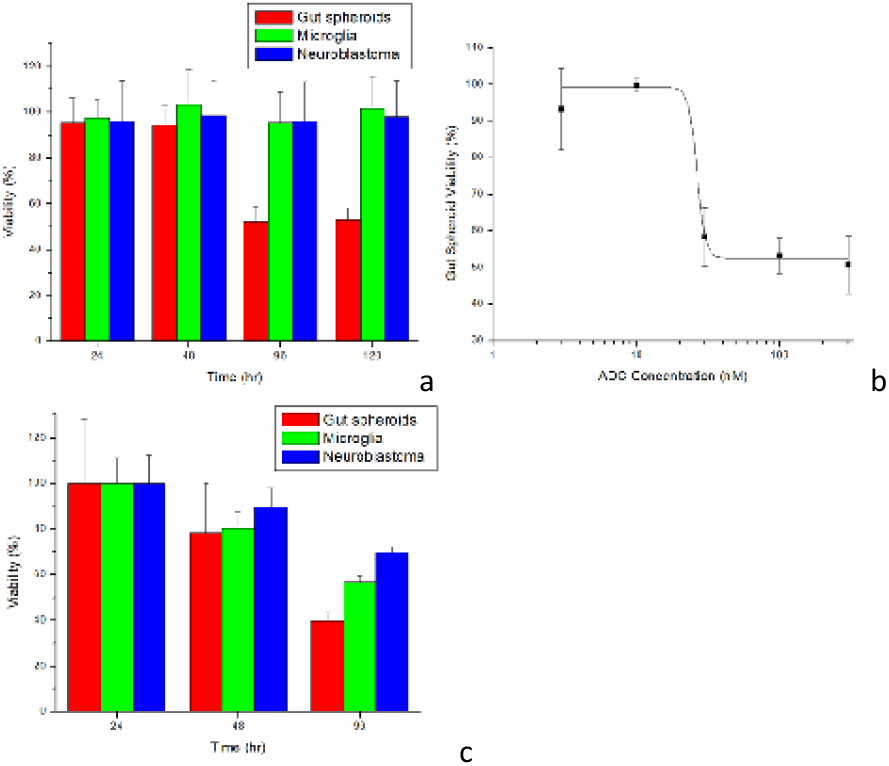
(a) Evaluation of on/off-target tumor killing kinetics using ADC Labetuzumab Govitecan (b) Effect of ADC concentration on target cells’ viability after 120 hours of treatment. (c) Evaluation of on-target and off-target killing kinetics using by-stander effects.

## Discussion and Conclusions

The OrganRX™ platform presents a significant advancement in multiorgan systems, offering a physiologically relevant, gravity-driven, net-unidirectional recirculation system capable of modeling inter-organ interactions and systemic drug effects with high fidelity. CFD modeling and impedance-based flow validation confirm the platform’s ability to generate consistent, quantifiable shear profiles across organ compartments, supporting its application in preclinical pharmacokinetic and toxicity studies. By integrating multiple organ models, including gut, liver, kidney, brain, and endothelium, within a modular microfluidic plate with dynamic fluid shear, OrganRX™ enables the modeling of vascularization, and drug metabolism under tunable conditions. This work demonstrates the system’s functional utility through a series of complex biological assays, including 3D tissue construction, vascular formation, ROS quantification, senescence evaluation, and biochemical marker analysis. Albumin and urea measurements from liver–kidney co-cultures validate metabolic interactions across organ models, offering translational relevance to human physiology. The OrganRX™ platform enables physiologically relevant modeling of human immune-mediated tumor responses within an interconnected multi-organ microenvironment under dynamic recirculating flow conditions. Compared with conventional static culture systems, the flow-based organ model platform demonstrated sustained cytokine signaling, enhanced immune-mediated cytotoxicity, improved discrimination between matched and mismatched immune responses, and more robust T-cell activation. The system successfully modeled antigen-specific CD8^+^ T-cell therapies, bispecific antibody-mediated killing using cibisatamab, and ADC-mediated cytotoxicity using Labetuzumab Govitecan, while simultaneously evaluating on-target efficacy, off-target toxicity, and bystander effects across multiple tissues. These studies demonstrate the utility of the OrganRX™ platform as a human-relevant tool for immuno-oncology, biologics safety assessment, cytokine response analysis, and evaluation of complex immune-tissue interactions that are difficult to reproduce in conventional static culture systems. Although the present studies demonstrate the utility of OrganRX™ for evaluating immune-mediated efficacy and toxicity, additional benchmarking studies^83^ are needed to establish clinical predictivity. Future validation should assess whether the platform can reproduce known efficacy rank ordering, target-dependent activity, persistence, exhaustion, and toxicity profiles of approved and investigational cell therapies. Similarly, ADC validation studies should determine the ability of the system to capture target expression dependence, payload-specific mechanisms, bystander killing, therapeutic index differences, and clinically recognized organ toxicities. Successful back-translation of therapies with known clinical outcomes would provide strong evidence that multiorgan immune-competent platforms can serve as predictive tools for identifying safety liabilities, prioritizing candidates, and accelerating the development of safer and more effective immunotherapies.

In summary, the OrganRX™ platform, built on the FPGA technology, represents a significant advancement in MOOC research by addressing long standing challenges in physiological relevance, scalability, automation compatibility and modularity bridging the gap between conventional in vitro models and human biology. The studies presented here demonstrate the versatility and functionality of the platform for modeling inter-organ communication^84^, immune-mediated responses, neuroinflammation, senescence, and biologics-associated toxicity. Furthermore, compatibility with automated imaging, biochemical assays, and transcriptomic analysis positions OrganRX™ as a versatile platform for medium-to high-throughput evaluation of small molecules, biologics, and emerging immunotherapy applications^85–88^. While these studies establish the broad utility of OrganRX™ across multiple application areas, additional benchmarking studies against clinically characterized therapeutics and human outcomes will be required to establish predictive performance for specific drug modalities such as ADCs, bispecific antibodies, checkpoint inhibitors, and cell therapies. Future studies focused on benchmarking against clinically characterized therapeutics and human outcomes will further establish its utility as a scalable multiorgan human-relevant platform for drug discovery, safety assessment, disease modeling, and precision medicine applications^89^.

## Author contributions

Conceptualization and Funding acquisition was by JC, Data curation by HCW, CJC, Formal analysis by HCW, Investigation by HCW, CJC, JAJJ, INB, GK, Methodology by HCW, CJC, JAJJ, INB, GK, JK, HSS, Project administration by JC, Resources by JC, Software by AJC, Supervision by JC, Validation by HCW, CJC, JAJJ, INB, GK, JK, HSS, Visualization by HCW, CJC, Writing original draft CJC, Writing review and editing by JC, HCW, INB, JAJJ.

## Data availability

Z-stack images of selected organ models will be posted in the Supplemental Information Section. RTPCR Data will be made available. The protocol for multiorgan culture using 3D gel is available in mathematical formula-enabled forms on the Biopico Systems Inc. website: https://biopico.com/organ-culture-estimate/

## Acknowledgements

The authors acknowledge the funding from NIH (1R43ES032357-01, 1R44GM139413-01 and 1R43AG073040-01) and NASA (80NSSC24PB276) for the development of the multiorgan plate. Dr. Adeela Syed (UC Irvine) is acknowledged for helping to acquire images from LSM 780 confocal microscope at Optical Biology Core using linear unmixing. Dr. Jay Sharma (Celprogen), Dr. Masashi Kitazawa (UC Irvine), Dr. Siok Lam Lim (UC Irvine), Dr. Subramanian Veedamali (UC Irvine) and Mr. Trevor Teafatillar (UC Irvine) are acknowledged for helping with cell culture and performing preliminary experiments on the OrganRX system.

## Supplement Note

### Analytical Calculation of Flow Velocity and Shear Stress

Assuming laminar steady-state flow of media through the channel, the velocity profile is given by:

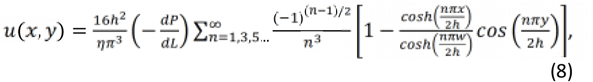

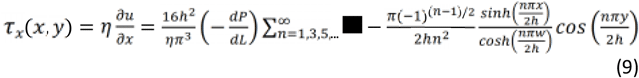

Where *h* = 0.7 mm is the half-height of the channel, *w* = 2 mm is the half-width of the channel, ™ *w* ≤ *x* ≤ *w*, and ™ *h* ≤ *y* ≤ *h*. The distribution of the shear stress components are given by:

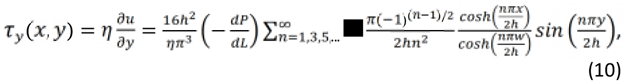

Where *η* = 1 cP is the viscosity of the media, which is modeled as water. We evaluated the shear stresses at *x* =*w* and *y* = *h* to find the maximum values. The average wall shear stress can also be calculated^90^ using integral means:

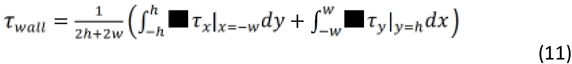

The volumetric flow rate through the channel is given by:

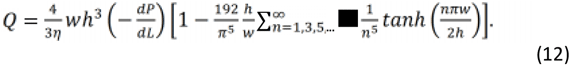

From this, the average velocity magnitude (speed) can be computed by:

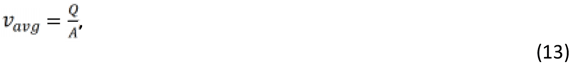

Where *A* =4*wh* is the cross-sectional area of the rectangular channel^91^. The equations for the velocity field (Equation 8) and the subsequent shear stresses (Equations 9 and 10) are valid for steady-state conditions.

Images embedded in text

